# Genetic Association of Arterial Stiffness Index with Blood Pressure and Coronary Artery Disease

**DOI:** 10.1101/453878

**Authors:** Seyedeh M. Zekavat, Mary Haas, Krishna Aragam, Connor Emdin, Amit V. Khera, Derek Klarin, Hongyu Zhao, Pradeep Natarajan

**Affiliations:** Program in Medical and Population Genetics, Broad Institute of MIT and Harvard, Cambridge, MA; Yale School of Medicine, New Haven, CT; Computational Biology & Bioinformatics Program, Yale University, New Haven, CT; Center for Genomic Medicine, Massachusetts General Hospital, Boston, MA; Harvard Medical School, Boston, MA; Cardiovascular Research Center, Massachusetts General Hospital, Boston, MA; Department of Biostatistics, Yale School of Public Health, New Haven, CT

**Keywords:** Arterial Stiffness, Blood Pressure, Coronary Artery Disease, Genetic Epidemiology, Mendelian Randomization, Population Genetics

## Abstract

**Background:** Arterial stiffness index (ASI) is independently associated with blood pressure and coronary artery disease (CAD) in epidemiologic studies. However, it is unknown whether these associations represent causal relationships.

**Objectives:** Here, we assess whether genetic predisposition to increased ASI is associated with elevated blood pressure and CAD risk.

**Methods:** Genome-wide association analysis (GWAS) of finger photoplethysmography-derived ASI was performed in 131,686 participants from the UK Biobank. Across UK Biobank participants not in the ASI GWAS, a 6-variant ASI polygenic risk score was calculated. The ASI polygenic score was associated with systolic and diastolic blood pressures (SBP, DBP, N=208,897), and with incident CAD over 10 years follow-up (N=223,061; 7,534 cases). The lack of CAD association observed was replicated among 184,305 participants (60,810 cases) from the Coronary Artery Disease Genetics Consortium (CARDIOGRAMplusC4D).

**Results:** We replicated prior reports of the epidemiologic association of ASI with SBP (Beta 0.55mmHg, [95% CI, 0.45–0.65], *P*=5.77×10^−24^), DBP (Beta 1.05mmHg, [95% CI, 0.99–1.11], *P*=7.27×10^−272^), and incident CAD (HR 1.08 [95% CI, 1.04–1.11], *P*=1.5×10^−6^) in multivariable models. While each SD increase in genetic predisposition to elevated ASI was highly associated with SBP (Beta 4.63 mmHg [95% CI, 2.1–7.2]; *P*=3.37×10^−4^), and DBP (Beta 2.61 mmHg [95% CI, 1.2–4.0]; *P*=2.85×10^−4^), no association was observed with incident CAD in UK Biobank (HR 1.12 [95% CI, 0.55–2.3]; *P*=0.75), or with prevalent CAD in CARDIOGRAMplusC4D (OR 0.56 [95% CI, 0.26–1.24]; *P*=0.15).

**Conclusions:** A genetic predisposition to higher ASI was associated with elevated blood pressure but not with increased risk of developing CAD.

**Condensed Abstract:** Arterial stiffness index (ASI) is proposed by some as a surrogate of blood pressure and coronary artery disease (CAD) risk based on epidemiologic analyses. We tested whether genetic predisposition to increased ASI is associated with elevated blood pressure and CAD risk to assess whether these represent causal relationships. We find that a genetic predisposition to higher ASI is associated with elevated systolic (Beta 4.63 mmHg [95% CI, 2.1–7.2]) and diastolic blood pressures (Beta 2.61 mmHg [95% CI, 1.2–4.0]) in the UK Biobank, but not associated with incident CAD in the UK Biobank (*P*=0.75) or with prevalent CAD in CARDIOGRAMplusC4D (*P*=0.15). These data support a causal relationship of ASI with blood pressure but do not support the notion that ASI is a suitable surrogate for CAD risk.

## Introduction

Arterial stiffness index (ASI), as measured non-invasively via pulse wave analysis, is independently associated with cardiovascular disease risk in multiple epidemiological studies (1–9). Increased vascular resistance and diminished viscoelasticity are key features of vascular aging which were previously associated with systolic hypertension (5), coronary artery disease (CAD) (2,4,7), and all-cause mortality (10). Arterial stiffness may be influenced by variations in collagen, elastin, smooth muscle tone, and endothelial dysfunction, in addition to other factors (11–17). Carotid-femoral (aortic) pulse wave velocity is the ‘gold-standard’ approach for assessing arterial stiffness. ASI measurement using finger infrared analysis is a scalable, non-invasive approach to assess ASI and is correlated with aortic pulse wave velocity (18–20).

While arterial stiffness measures are associated with cardiovascular diseases (1–8), whether the associations are causal is not clear. For example, non-causal risk factors, such as high-density lipoprotein cholesterol for CAD, are good risk predictors but are disappointing therapeutic targets (21–26). Lifestyle factors are separately linked to arterial stiffness and cardiovascular diseases, potentially confounding the observed relationships (27). Furthermore, reverse causality could lead to a statistically robust but non-causal relationship. For example, individuals with increased arterial stiffness might develop cardiovascular disease because of reduced exercise (28).

Some propose that ASI should be considered a non-invasive surrogate end point for cardiovascular events largely based on robust epidemiological associations (29–31)(2,32–38). Understanding whether ASI causally mediates CAD, independent of blood pressure, may help determine whether ASI is a suitable surrogate end point for CAD separate from its utility as a risk predictor. Mendelian randomization uses human genetics for causal inference by leveraging the random assortment of genetic variants during meiosis at conception, which diminishes susceptibility to confounding or reverse causality (39). Here, we used Mendelian randomization to determine whether a genetic predisposition to increased ASI is associated with elevated blood pressure and increased risk for incident CAD.

## Methods

### UK Biobank study participants and phenotypes

Individual-level genomic data and longitudinal phenotypic data from the UK Biobank, a large-scale population-based dataset consisting of genotype and phenotype data in approximately 500,000 volunteer participants collected from 2007–2017, was used.

Clinical disease definitions are detailed in **Supplementary Table 1**. In summary, the main outcome, CAD, was defined by billing codes for heart attack, angina pectoris, unstable angina, myocardial infarction, coronary atherosclerosis, coronary artery revascularization, and other acute, subacute, and chronic forms of ischemic heart disease, or with self-reported angina, heart attack/myocardial infarction, coronary angioplasty +/- stent, or coronary artery bypass graft (CABG) surgery. We also assessed systolic and diastolic blood pressures, and adjusted for blood pressure medications by adding 15 and 10 mmHg to systolic and diastolic blood pressures, respectively (40).

### Arterial stiffness index measurement

ASI was previously measured in the UK Biobank using the PulseTrace PCA2 (CareFusion, San Diego, CA), which uses finger photoplethysmography over a 10- to15-second timeframe to obtain the pulse waveform from an infrared sensor clipped to the end of the index finger. ASI (in m/s) was calculated by dividing standing height by the time between the systolic and diastolic peaks of the pulse waveform. ASI by this approach was previously correlated with aortic pulse wave velocity, which is regarded as the gold standard (18). ASI was inverse rank normalized for analysis (with mean = 0, SD = 1).

### Genotyping and imputation

Genome-wide genotyping was previously performed in the UK Biobank using two genotyping arrays sharing 95% of marker content: Applied Biosystems UK BiLEVE Axiom Array (807,411 markers in 49,950 participants) and Applied Biosystems UK Biobank Axiom Array (825,927 markers in 438,427 participants) both by Affymetrix (Santa Clara, CA) (41). Variants used in the present analysis include those also imputed using the Haplotype Reference Consortium reference panel of up to 39M SNPs (42,43).

### Quality control and variant annotation

Poor quality variants and genotypes were filtered as previously described (41). We further filtered out individuals from both genetic and epidemiological analyses using the following genetic criteria: non-white or not of British ancestry, gender mismatch between reported and genotypic genders, sex chromosome aneuploidy, or one from each pair of 1^st^ or 2^nd^ degree relatives (**Supplementary Table 2**). Non-consenting individuals with prevalent peripheral arterial disease, aortic valve disease, or CAD were excluded, as were extreme outliers for any of the arterial pulse wave phenotypes listed in **Supplementary Table 3**. Extreme outliers were determined by adjusting the traditional box and whisker upper and lower bounds and accounting for skewness in the phenotypic data identified using the Robustbase package in R (setting range=3) (https://cran.r-project.org/web/packages/robustbase/robustbase.pdf).

After filtering samples, variants were further filtered by the following criteria: not in Hardy-Weinberg Equilibrium (*P*<1×10^−10^), low imputation quality (INFO score < 0.3), call rate < 95%, and minor allele frequency < 0.05% (minor allele count < 66).

Variant consequences were annotated using with Ensembl’s Variant Effect Predictor (VEP), ascribing the most severe consequence and associated gene among the canonical transcripts present for each variant^(44)^. The Hail v0.1 software (https://hail.is) was used to perform quality control and variant annotation (45).

### Epidemiological association analyses with arterial stiffness index

Epidemiological association of ASI with blood pressure phenotypes and incident CAD was performed using linear regression and Cox proportional hazards model, respectively, in R (version 3.3, R Foundation, Vienna, Austria). For CAD, adjustment was performed for age, sex, ever smoking status, heart rate at pulse wave analysis, prevalent hypertension, prevalent hypercholesterolemia, prevalent diabetes, alcohol intake (self-reported alcohol intake of at least once per month), exercise (self-reported exercise of at least 3x per week), and vegetable intake (self-reported intake of at least 6 tablespoons of vegetable intake per day). The same adjustment variables were used for SBP and DBP, except prevalent hypertension was not included as a covariate.

Analyses were performed using a Cox proportional hazards model for incident CAD, and linear regression for the blood pressure traits. The threshold for significance for the three primary phenotypes was assigned as alpha = 0.05/3 tests = 0.017.

### Genome-wide association analysis of arterial stiffness

A genome-wide association of ASI was performed using individual-level data from 131,686 individuals of European descent from the UK Biobank, collected from 2007 to 2017. Each variant was individually associated with ASI in an additive linear regression model and adjusted for sex, age, smoking status, genotyping array type, and the first ten principal components of ancestry (46). Only variants with minor allele frequency > 0.05% (minor allele count > 66) were considered. *P* < 5×10^−8^ was considered to be significant. The Hail software version 0.1 (https://hail.is) was used for genome-wide association analysis (45).

### Mendelian randomization

An additive genetic risk score (GRS) was calculated as 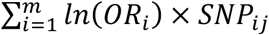 were *m* is the number of SNPs, *ln*(*OR_i_*) is the weight for *SNP_i_* from the discovery sample, *SNP_ij_* is the number of alleles (i.e., 0, 1, or 2) for *SNP_i_* in person *j* in the validation sample. Six independent variants (linkage disequilibrium r^2^ < 0.25 within 500kb windows) demonstrating at least suggestive association with ASI (*P*<5×10^−7^) were included in the GRS. The raw GRS was calculated for each individual using PLINK (47), inverse rank normalized, then re-scaled such that one unit increase in the GRS was equivalent to a one standard deviation (SD) increase in ASI.

To confirm that the GRS for ASI was a strong instrument for ASI, an F-statistic for the instrument was calculated in the UK Biobank. An F-statistic is a measure of the significance of an instrument (the GRS) for prediction of the exposure (ASI), controlling for additional covariates (age, sex, ever smoked, 10 principal components of ancestry, and genotyping array type). An F-statistic greater than 10 is evidence of a strong instrument. Furthermore, sensitivity analyses were performed to evaluate for associations between the ASI GRS and potential environmental confounders including sex, ever smoking status, diet (alcohol intake, vegetable intake), and exercise frequency among individuals not in the ASI genome-wide association analyses.

A linear regression model was used to associate the ASI GRS with systolic and diastolic blood pressures. A Cox proportional hazards model was used to associate ASI GRS with incident CAD. For CAD, adjustment was performed for age, sex, ever smoking status, heart rate at blood pressure measurement, prevalent hypertension, prevalent hypercholesterolemia, prevalent diabetes, alcohol intake (self-reported alcohol intake of at least once per month), exercise (self-reported exercise of at least 3x per week), and vegetable intake (self-reported intake of at least 6 tablespoons of vegetable intake per day), where indicated. The same adjustment variables were used for SBP and DBP, except for prevalent hypertension.

### 2-Sample Mendelian randomization with coronary artery disease

To address potential power limitations from the lack of association between ASI and CAD, we also pursued 2-sample Mendelian randomization using variant-level summary statistics from prior genome-wide association analyses of CAD from several independent case-control studies, specifically 184,305 individuals from the Coronary Artery Disease Genetics Consortium (CARDIOGRAMplusC4D) (48). The effect estimates and standard errors for the six GRS variants for ASI (from UK Biobank) and for CAD (from CARDIOGramplusC4D) were used to perform robust, penalized inverse variance weighted (IVW) 2-sample Mendelian randomization using the MendelianRandomization package in R (49,50). IVW 2-sample Mendelian randomization uses a weighted linear regression of the ratio of the SNP effects on the outcomes to the SNP effects on the risk factor, without using an intercept term. The threshold for significance was defined as alpha = 0.05.

Additionally, analyses were performed to evaluate the reverse association, of CAD causally impacting ASI. 77 known, independent, genome-wide significant CAD locus variants were identified across several published sources (48,51–53) (**Supplementary Table 9**). These 77 CAD locus variants were used as an instrument in 2-sample Mendelian randomization to evaluate whether CAD causally affects ASI.

## Results

### Baseline characteristics

A total of 131,686 individuals in the UK Biobank had ASI measured, genotype data available, and passed quality control (**Supplementary Table 2**). Among these individuals, median age was 59 (IQR 51–63) years, 53.8% were female, 4.6% had diabetes, 27.1% had hypertension, and 12.9% had hypercholesterolemia. Median SBP was 139 (IQR 127–153) mmHg, median DBP was 82 (IQR 75–89) mmHg. 44.1% of individuals were prior or current smokers, and 10.1% of individuals were on antihypertensive medications (**Table 1**). The median ASI was 9 (IQR 7–11) m/s (**Supplementary Table 3**).

**Table 1:**
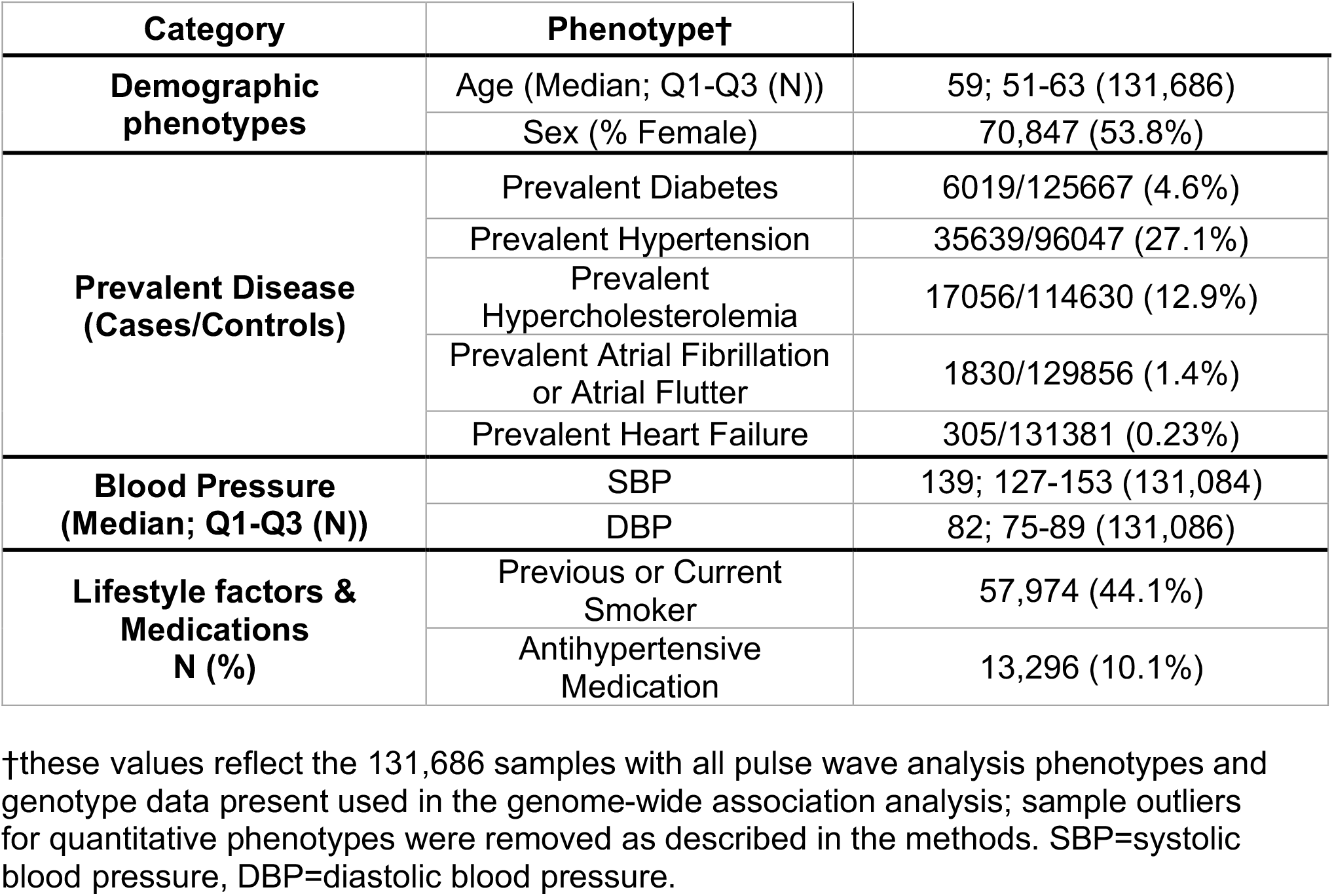
Baseline characteristics of analyzed participants with arterial stiffness index and genotypes.

| Category | Phenotype† |  |
| --- | --- | --- |
| **Demographic phenotypes** | Age (Median; Q1-Q3 (N)) | 59; 51-63 (131,686) |
|  | Sex (% Female) | 70,847 (53.8%) |
| **Prevalent Disease (Cases/Controls)** | Prevalent Diabetes | 6019/125667 (4.6%) |
|  | Prevalent Hypertension | 35639/96047 (27.1%) |
|  | Prevalent Hypercholesterolemia | 17056/114630 (12.9%) |
|  | Prevalent Atrial Fibrillation or Atrial Flutter | 1830/129856 (1.4%) |
|  | Prevalent Heart Failure | 305/131381 (0.23%) |
| **Blood Pressure (Median; Q1-Q3 (N))** | SBP | 139; 127-153 (131,084) |
|  | DBP | 82; 75-89 (131,086) |
| **Lifestyle factors & Medications N (%)** | Previous or Current Smoker | 57,974 (44.1%) |
|  | Antihypertensive Medication | 13,296 (10.1%) |

†these values reflect the 131,686 samples with all pulse wave analysis phenotypes and genotype data present used in the genome-wide association analysis; sample outliers for quantitative phenotypes were removed as described in the methods. SBP=systolic blood pressure, DBP=diastolic blood pressure.

### Epidemiological associations of ASI

Univariate association of cardiovascular risk factors with ASI showed the following associations with at least nominal significance (*P*<0.05): for age (0.024 SD/year, *P*<1×10^−300^), sex (0.40 SD higher in males, *P*<1×10^−300^), blood pressure medication (0.34 SD, *P*=1.4×10^−317^), hypertension (0.21 SD, *P*=1.4×10^−269^), hypercholesterolemia (0.20 SD, *P*=4.1×10^−137^), diabetes (0.20 SD, *P*=9.1×10^−54^), ever smoking (0.18 SD, *P*=3.0×10^−250^), exercise ≥3x/wk (−0.16 SD, *P*=2.9×10^−66^), alcohol intake ≥1x/mo (0.05 SD, *P*=3.3×10^−20^), and ≥6 tablespoons vegetable intake per day (−0.063 SD, *P*=3.1×10^−4^) (**Supplementary Table 4**).

For the associations of ASI with SBP and DBP, both univariable and multivariable, adjusting for age, sex, smoking status, prevalent hypercholesterolemia, prevalent diabetes, vegetable intake, alcohol intake, and exercise, analyses showed strong associations (**Figure 1A**). Each SD of ASI was associated with elevated SBP by 0.55 mmHg ([95% CI, 0.45–0.65], *P*=5.77×10^−24^) and DBP by 1.05 mmHg ([95% CI, 0.99–1.11], *P*=7.27×10^−272^).

ASI was also significantly independently associated with incident CAD, adjusting for age, sex, ever smoking status, heart rate, prevalent hypertension, prevalent hypercholesterolemia, prevalent diabetes, vegetable intake, alcohol intake, and exercise (HR 1.08 per SD ASI [95% CI, 1.04–1.11], *P*=7.67×10^−6^) (**Figure 2A**).

**Figure 1:**
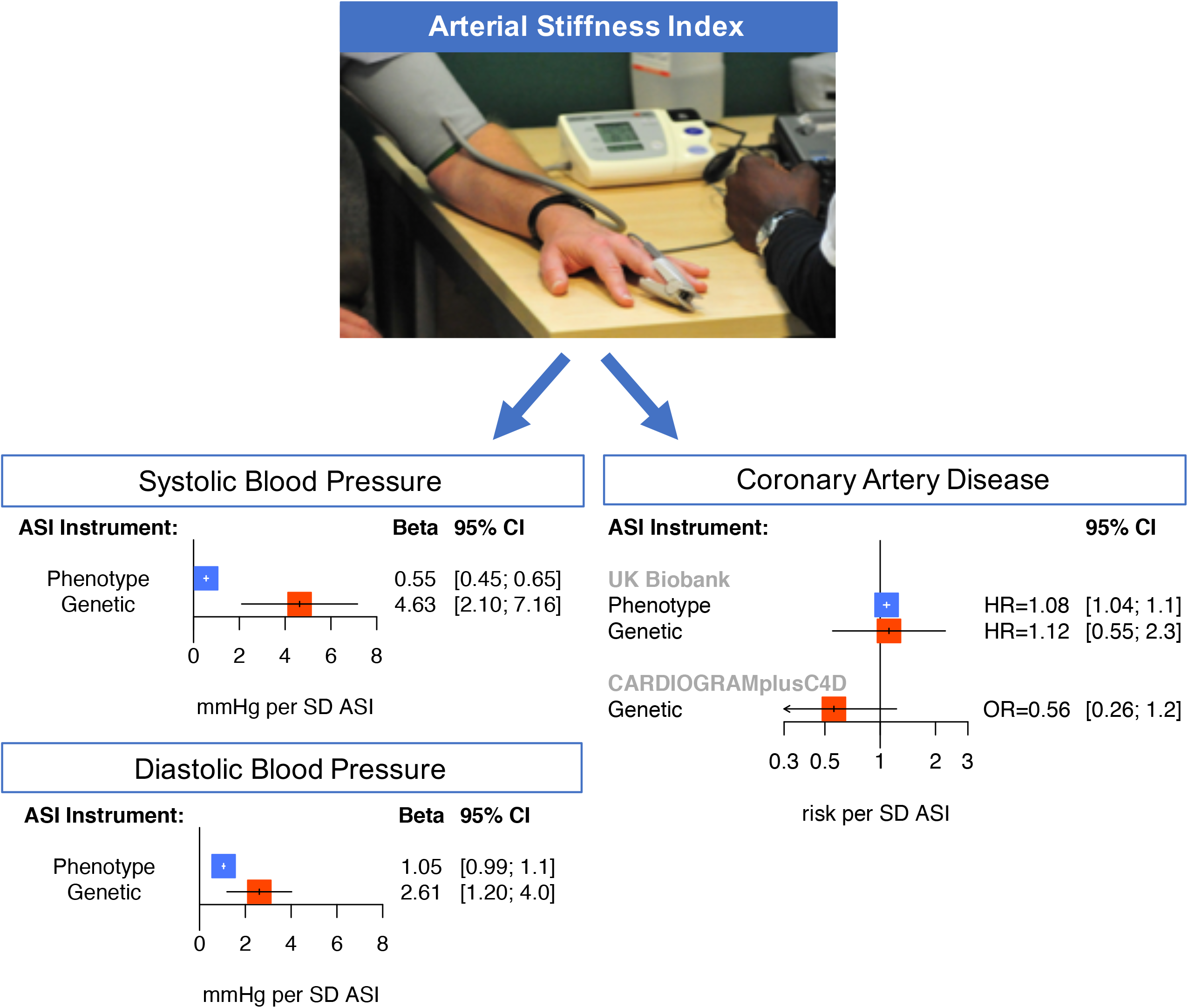
Epidemiologic and genetic associations of arterial stiffness index with blood pressure. Association between (A) phenotypic ASI, and, (B) genotypic ASI (ie: the ASI GRS), with systolic and diastolic blood pressures in the UK Biobank. Results are presented as both unadjusted and, separately, adjusted by age, sex, smoking status, prevalent hypercholesterolemia, prevalent diabetes, heart rate, vegetable intake, alcohol intake, and exercise frequency. For the ASI GRS instrument, analysis was performed excluding individuals used in the ASI genome-wide association study. Effect estimates represent mmHg increase in blood pressure resulting from (A) 1 SD increase in ASI phenotype, and (B) 1SD increase in genetically-mediated ASI from the ASI GRS.
ASI = Arterial stiffness index, DBP = diastolic blood pressure, GRS = genetic risk score, SBP = systolic blood pressure, SD = standard deviation

### Genome-wide association analysis of ASI

A genome-wide association analysis of ASI was performed among 131,686 individuals and 13,995,214 variants in the UK Biobank. A quantile-quantile plot of the genome-wide association statistics did not show substantial genomic inflation (*λ* = 1.05) (**Supplementary Figure 1**). Two genome-wide significant loci were identified (*P*<5×10^−8^), the top variants of which were in second intron of *TEX41* (rs1006923, −0.025 SD, *P*=3.7×10^−10^, minor allele frequency (MAF)=0.32), and first intron of *FOXO1* (rs7331212, −0.024 SD, *P*=9.3×10^−9^, MAF=0.26). Three additional suggestive loci (P<5×10^−7^) were also identified, of which the top variants are intronic variants in *COL4A2* (rs872588, - 0.020 SD, *P*=2.3×10^−7^, MAF=0.42), *RNF126* (rs1009628, −0.027 SD, *P*=1.2×10^−7^, MAF=0.15), and *TCF20* (rs55906806, −0.024 SD, *P*=2.4×10^−7^, MAF=0.20). Through chromatin conformational changes, intronic variants at *TEX41* and *COL4A2* may influence gene expression at nearby enhancers Supplementary Results, **Supplementary Figure 2**). Interrogation of disruptive protein-coding variants yielded moderate association for *HFE* p.Cys282Tyr (MAF 0.076), the most common variant implicated in hereditary hemochromatosis (**Supplementary Results, Supplementary Table 4**).

### Mendelian randomization in the UK Biobank

Six independent and at least suggestive (*P*<5×10^−7^) variants were used towards an ASI genetic risk score (GRS) (**Supplementary Table 5**). The raw ASI GRS was associated with a 0.85 SD increase in ASI (SE: 0.072; *P*=8.0×10^−32^). The F-statistic of the GRS was 123 (recommended F-statistic > 10), suggesting high instrument strength. The inverse-rank normalized GRS was re-scaled such that each unit reflected one SD in ASI for comparison with the phenotypic associations (**Supplementary Figure 3**). Sensitivity analysis was performed to evaluate for associations between the ASI GRS and potential environmental confounders including sex, ever smoking status, diet (alcohol intake, vegetable intake), and exercise frequency. No significant associations between the ASI GRS and environmental confounders were observed (**Supplementary Table 6**).

A 1-SD increase in genetically-mediated ASI was significantly associated with elevated SBP (Beta 4.63 mmHg [95% CI, 2.1–7.2]; *P*=3.37×10^−4^), and DBP (Beta 2.61 mmHg [95% CI, 1.2–4.0]; *P*=.85×10^−4^), independent of cardiometabolic risk factors (age, sex, and smoking status, prevalent hypercholesterolemia, prevalent diabetes, heart rate, vegetable intake, alcohol intake, and exercise frequency) (Figure 1B).

The ASI GRS, however, was not associated with incident CAD in UK Biobank in an unadjusted model (HR 1.3 [95% CI, 0.64–2.6]; *P*=0.47) or an adjusted model including age, sex, smoking status, prevalent hypertension, prevalent hypercholesterolemia, prevalent diabetes, heart rate, vegetable intake, alcohol intake, and exercise frequency as covariates (HR 1.12 [95% CI, 0.55–2.3]; *P*=0.75) (Figure 2B).

**Figure 2:**
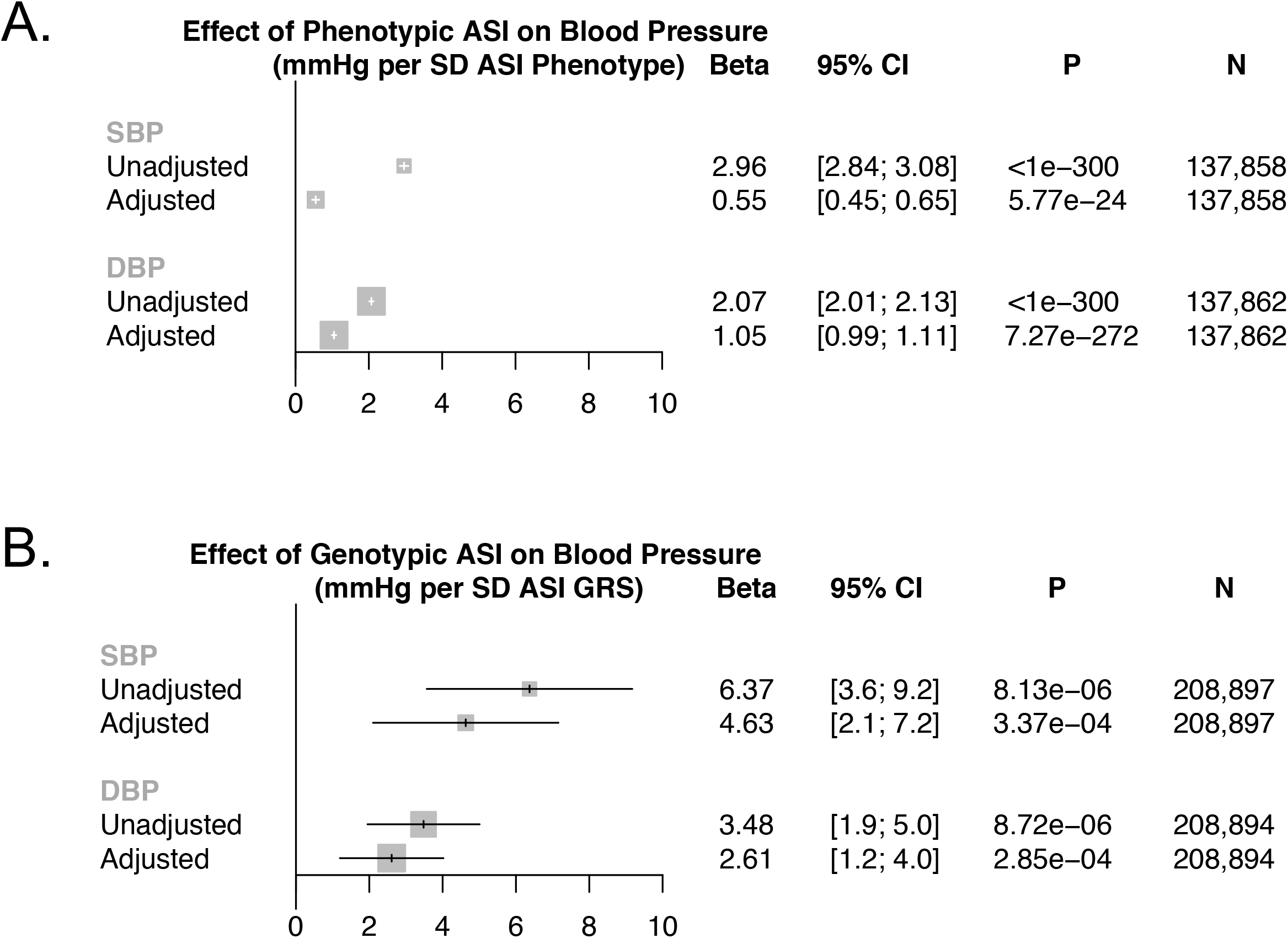
Epidemiologic and genetic associations of arterial stiffness index with incident coronary artery disease. Association between (A) phenotypic ASI, and, (B) the ASI GRS, with incident coronary artery disease in the UK Biobank. Results are presented as both unadjusted (cyan) and adjusted (purple) by age, sex, smoking status, prevalent hypertension, prevalent hypercholesterolemia, prevalent diabetes, heart rate, vegetable intake, alcohol intake, and exercise frequency. For the ASI GRS instrument, analysis was performed excluding individuals used in the ASI genome-wide association study. Hazard ratios represent increased risk of incident CAD resulting from (A) 1 SD increase in ASI phenotype, and (B) 1SD increase in genetically-mediated ASI from the ASI GRS. Sample sizes for (A) the phenotypic association are 3,692 cases, 126,615 controls, and for (B) the genotypic association are 7,534 cases, 215,527 controls.
ASI = Arterial stiffness index, CAD = coronary artery disease, GRS = genetic risk score, HR = hazard ratio, SD = standard deviation.

### 2-Sample Mendelian randomization with coronary artery disease

To address potential power limitations impeding association of ASI GRS with incident CAD in the UK Biobank, we also pursued 2-sample Mendelian randomization between ASI and prevalent CAD using variant-level summary statistics from 184,305 separate individuals in the Coronary Artery Disease Genetics Consortium (CARDIOGRAMplusC4D) (48). Robust, penalized inverse-variance weighted 2-sample Mendelian randomization similarly did not demonstrate an association between genetically-elevated ASI and CAD (OR 0.56 [95% CI, 0.26–1.24], *P*=0.15) (Figure 3). Furthermore, the six variants showing suggestive association with ASI did not demonstrate a significant positive association with CAD across several different 2-sample Mendelian randomization methods (**Supplementary Table 8**).

**Figure 3:**
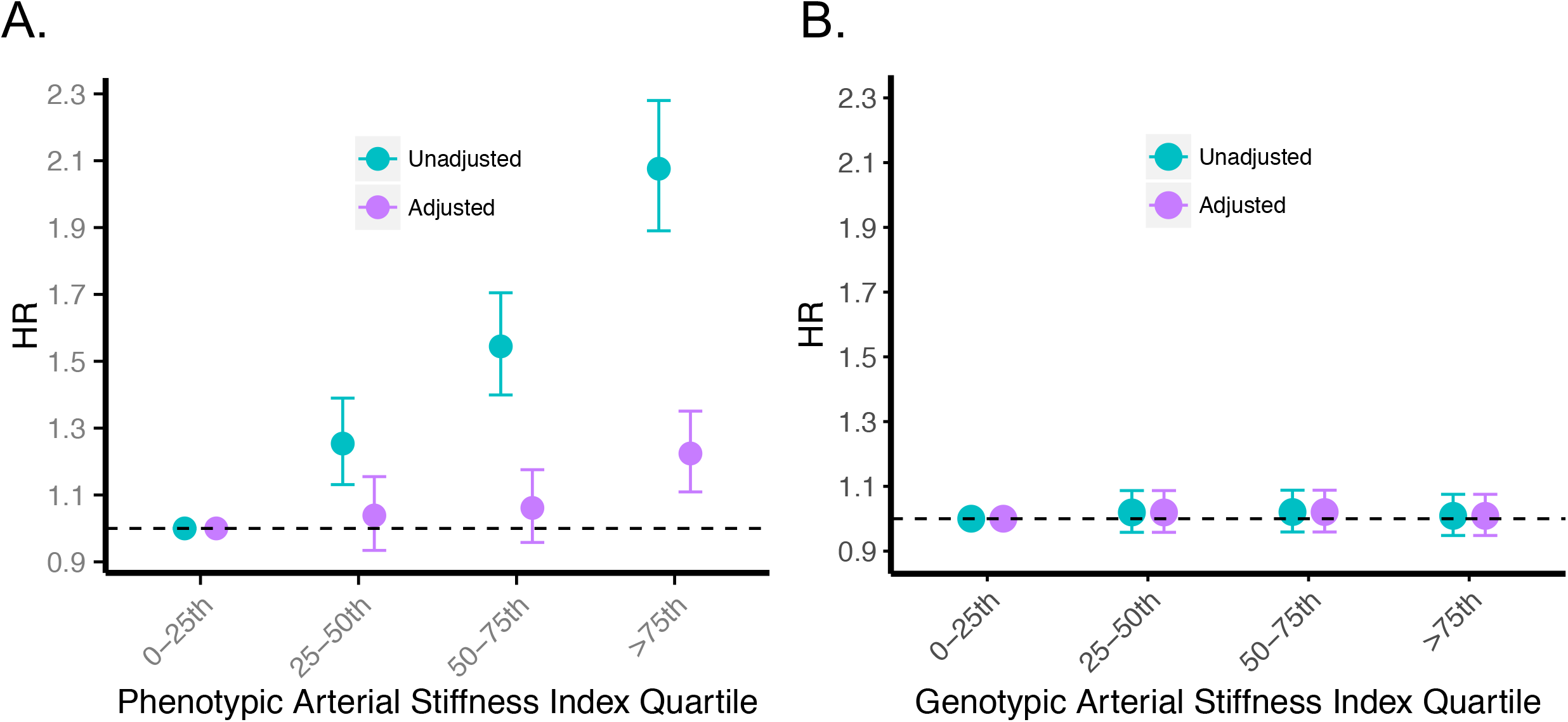
One-and two-sample Mendelian randomization analyses of arterial stiffness index with coronary artery disease. Association between the ASI GRS and incident CAD in the UK Biobank, as well as prevalent CAD in the CARDIOGRAMplusC4D consortium. Incident CAD results were derived using individual-level data from the UK Biobank and adjusting by cardiometabolic risk factors (age, sex, smoking status, prevalent hypertension, prevalent hypercholesterolemia, prevalent diabetes, heart rate, vegetable intake, alcohol intake, and exercise frequency). Individuals used in the ASI genome-wide association study were excluded in the analyses. Prevalent CAD results were derived from summary-level genome-wide association data from the CARDIOGRAMplusC4D consortium using robust, penalized inverse-variance weighted 2-sample Mendelian randomization. For the ASI GRS instrument, analysis was performed excluding individuals used in the ASI genome-wide association study.
ASI = Arterial stiffness index, CAD = coronary artery disease, GRS = genetic risk score, HR = hazard ratio, OR = odds ratio

We also developed an expanded ASI polygenic score using 321 independent variants (P<1×10^−4^, LD r^2^< 0.25) to capture additional genetic variation of ASI. The expanded ASI polygenic score explained 3.3% of ASI variance conferring >80% power to detect the CAD effect estimate observed in epidemiologic analyses (i.e., OR=1.08) with alpha =0.05. With this approach, we again confirmed no significant association in inverse-variance weighted 2-sample Mendelian randomization (OR 0.95 [95% CI, 0.89–1.02], *P*=0.13).

77 genome-wide significant CAD loci from prior GWAS (48,52,53) were identified, and CAD risk effect estimates prior studies and ASI effect estimates from this study were catalogued (**Figure 4**). While 3 of 77 previously-associated CAD loci showed evidence of association with ASI (*P*<0.05/77=6.5×10^−4^), effect directions were inconsistent between ASI and CAD. For example, the variant rs9349379-A, an intronic variant in *PHACTR1*, was associated with increased ASI (0.015 SD, *P*=4.5×10^−5^) but decreased risk for CAD (OR= 0.87, *P*=1.8×10^−42^). Similarly, ASI-raising alleles at the *ZEB2-TEX41* and *ABO* loci decrease CAD risk, while ASI-raising alleles at *CYP17A1-CNNM2-NT4C2* and *SH2B3* increase CAD risk. Detailed variant-level summary statistics for these 77 CAD locus variants are provided in **Supplementary Tables 9–10**. These 77 CAD locus variants were also used as an instrument in 2-sample Mendelian randomization for a putative reverse association – whether a genetic susceptibility to CAD increases ASI.
No significant associations were observed across various 2-sample Mendelian randomization methods for the reverse association (**Supplementary Table 11**).

**Figure 4:**
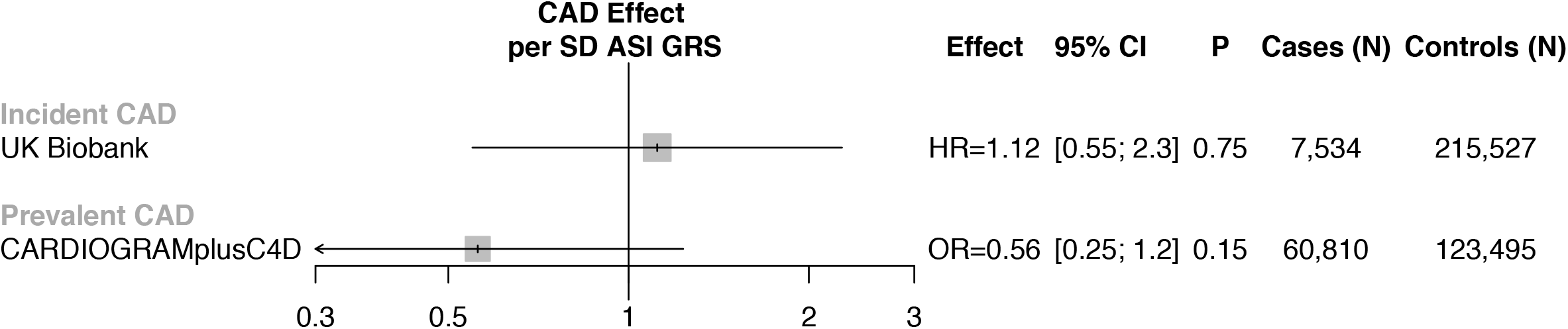
Comparison of variant level-effects with arterial stiffness index and with coronary artery disease showsinconsistency. Variant-level effect estimates (from CARDIOGRAMplusC4D) from variants at 77 independent known CAD loci, were compared to their ASI associations. Highlighted are 5 out of the 77 variants with at least suggestive significance with ASI (P<0.005), showing that ASI-raising alleles have inconsistent effects on CAD risk. The variant-level summary statistics for these 77 variants across are detailed in Supplementary Tables 9–10.
ASI = arterial stiffness index, CAD = coronary artery disease

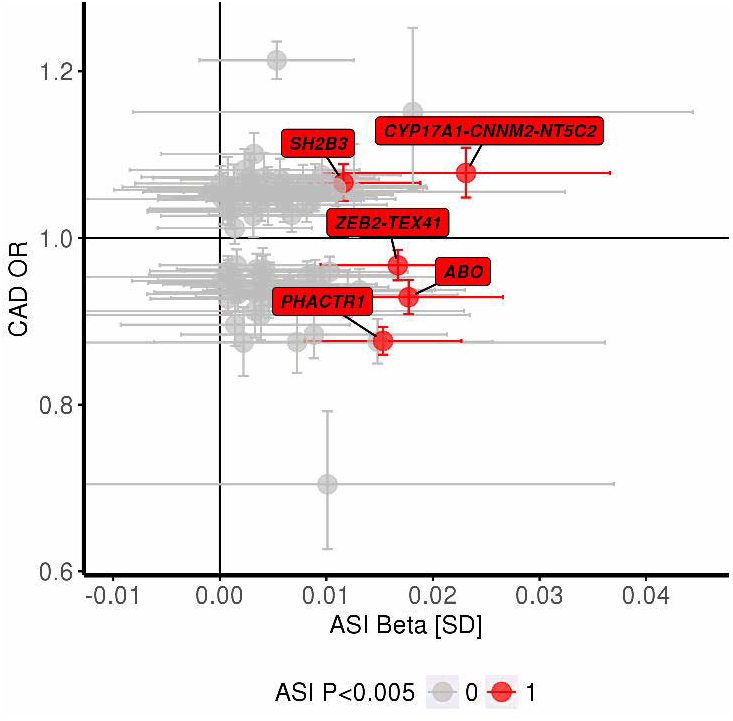

## Discussion

We performed the largest genome-wide association analysis to-date of a measure of vascular aging, ASI, in 131,686 individuals, and leveraged these observations to perform causal inference analyses with blood pressure and risk of CAD in up to 407,366 separate individuals. In our genome-wide association analyses, we discover the first genome-wide variants associated with ASI. We replicate the epidemiologic associations of ASI with blood pressure and CAD, and find that genetic analyses do indeed support a causal relationship between ASI and blood pressure. However, our genetic analyses do not support a causal relationship between ASI and CAD.

These results permit several conclusions. First, we observe strong epidemiologic and genetic association between ASI and blood pressure. These data indicate that non-invasive photoplethysmography, employed by a finger probe or potentially commercially-available wearable monitors that measure heart rate (54), may be used to impute continuous blood pressure, and that changes will track with blood pressure changes. However, given independent clinical effects and imperfect correlation, ASI measurement may complement blood pressure assessments. Second, there is a long-standing debate whether ASI precedes elevated blood pressure or vice versa (55). Compared to its phenotypic effect, the effect conferred by genetically-elevated ASI is 8.4-fold higher for SBP (4.63 mmHg for ASI GRS versus 0.55 mmHg for ASI phenotype) and 2.5-fold higher for DBP (2.61 mmHg for ASI GRS versus 1.05 mmHg for ASI phenotype), potentially representing the effects of life-long exposure to elevated arterial stiffness on blood pressure. This supports the notion that arterial stiffness may predate the onset of elevated blood pressure indicating that ASI may identify individuals at heightened risk for future blood pressure elevations.

Third, our epidemiological and genetic analyses indicate that ASI is an independent, non-causal risk factor for CAD. Arterial stiffness may be a parallel disrupted pathway in the setting of CAD, as opposed to an upstream causal mediating factor. Thus, while ASI monitoring may still serve as a good proxy for blood pressure, therapeutic modulation of ASI in isolation may not have a meaningful impact on CAD outcomes. Similarly, a recent study of twins showed that while carotid-femoral pulse wave velocity was heritable, it did not associate with 5-year progression of carotid intima media thickness (56). The lack of significance between genetically-elevated ASI and CAD is also consistent with prior mixed results in experimental models. Fragmentation of elastin fibers and deposition of collagen fibers are features of vascular aging implicated in arterial stiffness (57). However, murine models lacking elastin do not have endothelial damage, thrombosis, or inflammation which typically occur with atherosclerosis (58).

Furthermore, we found that while few variants associated with CAD show apparent association with ASI, our data indicate that ASI may not be mediating the apparent CAD risk. We observed generally inconsistent genetic effects between ASI and CAD risk. In particular, an intronic variant within *PHACTR1* (rs9349379-A), which was recently shown to influence endothelin-1 expression in the vasculature, is associated with decreased risk for CAD (59), increased blood pressure (60), and increased ASI. For this variant, the divergent directionalities of effect on CAD and blood pressure may be due to the differential expression of *EDNRA* versus *EDNRB* in the coronary arteries compared to peripheral vasculature (59). Additionally, genetic variants disrupting nitric oxide signaling at the *NOS3* and *GUCY13* loci influence both blood pressure and risk of CAD (61–63). Notably, in our study, risk variants at these loci were not strongly associated with ASI. Extensive prior experimental work linked nitric oxide signaling and endothelin-1 with endothelial function and vascular tone (64–68). Our data suggests that increased risk of CAD through these pathways is unlikely to be through changes in photoplethymsography-detected ASI but potentially through alternative vascular mechanisms.

While our study has several strengths, some limitations should be considered. First, lack of ASI genetic risk score association with CAD may be due to limited statistical power. Our replication of the lack of association using 2-sample Mendelian randomization including with an expanded polygenic score, combined with our analysis showing inconsistent effects of individual variants between CAD and ASI suggests that this is less likely. Second, our imputation of untreated blood pressure among those with prescribed hypertensives assumes a homogenous blood pressure effect across the population. Without prescription data in the UK Biobank, we are unable to account for different medication regimens and adherence.

## Conclusion

A genetic predisposition to higher ASI was associated with increased blood pressure, but not increased risk of CAD. Our data support the conclusion that finger photoplethysmography-derived ASI is an independent, causal risk factor for blood pressure and an independent, non-causal risk factor for CAD.

## Clinical Perspectives

### Core Clinical Competencies

A genetic predisposition to higher ASI was associated with elevated blood pressure, but not with elevated risk for CAD.

### Translational Outlook

Further research should be conducted to determine whether photoplethysmography-derived ASI may be used in wearables as a continuous proxy for blood pressure phenotypes for prevention and monitoring.

### Translational Outlook

Further research is required to understand whether the novel genes identified and implicated in ASI are suitable novel targets for blood pressure-lowering.

## Central Illustration: Epidemiologic and genetic associations of arterial stiffness with blood pressure and coronary artery disease

Association between phenotypic ASI, and separately, genotypic ASI (from the ASI GRS), with systolic and diastolic blood pressures, as well as with incident and prevalent CAD. Blood pressure results are adjusted by age, sex, smoking status, prevalent hypercholesterolemia, prevalent diabetes, heart rate, vegetable intake, alcohol intake, and exercise frequency. Blood pressure effect estimates represent mmHg increase in blood pressure resulting from (A) 1 SD increase in ASI phenotype, and (B) 1SD increase in genetically-mediated ASI from the ASI GRS.

Incident CAD associations in the UK Biobank, as well as prevalent CAD associations using the CARDIOGRAMplusC4D consortium, with phenotypic ASI and the ASI GRS are provided. Incident CAD results were derived using individual-level data from the UK Biobank and adjusting by cardiometabolic risk factors (age, sex, smoking status, prevalent hypertension, prevalent hypercholesterolemia, prevalent diabetes, heart rate, vegetable intake, alcohol intake, and exercise frequency). Prevalent CAD results were derived from summary-level genome-wide association data from the CARDIOGRAMplusC4D consortium using robust, penalized inverse-variance weighted 2-sample Mendelian randomization.

## Acknowledgements

The authors would like to acknowledge and thank the participants and staff of the UK Biobank and of the CARDIOGramplusC4D consortium cohorts.

## Abbreviations

ASI: arterial stiffness index
CAD: coronary artery disease
CARDIOGRAMplusC4D: Coronary Artery Disease Genetics Consortium
DBP: diastolic blood pressure
GRS: genetic risk score
PPG: photoplethysmography
SBP: systolic blood pressure

